# Metabolic abnormalities in the bone marrow cells of young offspring born to obese mothers

**DOI:** 10.1101/2023.11.29.569274

**Authors:** Elysse Phillips, Yem Alharithi, Leena Kadam, Lisa M. Coussens, Sushil Kumar, Alina Maloyan

**Affiliations:** Knight Cardiovascular Institute, Oregon Health & Science University, Portland, OR, 97239; Department of OB/GYN, Oregon Health & Science University, Portland, OR, 97239; Department of Cell, Development and Cancer Biology, Knight Cancer Institute, Oregon Health & Science University, Portland, OR, 97239

**Author notes:** **Corresponding Authors:** Alina Maloyan, Ph.D., Knight Cardiovascular Institute, and Sushil Kumar, Ph.D., Knight Cancer Institute, Oregon Health and Science University, 3181 SW Sam Jackson Park Rd, Portland, Oregon, 97239. and. These authors contributed equally to the paper (Elysse Phillips, Yem Alharithi, Sushil Kumar, Alina Maloyan). **Lead Contact**: Alina Maloyan, Ph.D., Knight Cardiovascular Institute, Oregon Health and Science University, 3181 SW Sam Jackson Park Rd, Portland, Oregon, 97239.; twitter.com/@AMaloyan. Requests for resources and reagents should be directed to and will be fulfilled by the lead contact, Alina Maloyan.

## Abstract

Intrauterine metabolic reprogramming occurs in obese mothers during gestation, putting the offspring at high risk of developing obesity and associated metabolic disorders even before birth. We have generated a mouse model of maternal high-fat diet-induced obesity that recapitulates the metabolic changes seen in humans. Here, we profiled and compared the metabolic characteristics of bone marrow cells of newly weaned 3-week-old offspring of dams fed either a high-fat (Off-HFD) or a regular diet (Off-RD). We utilized a state-of-the-art targeted metabolomics approach coupled with a Seahorse metabolic analyzer. We revealed significant metabolic perturbation in the offspring of HFD-fed vs. RD-fed dams, including utilization of glucose primarily via oxidative phosphorylation, and reduction in levels of amino acids, a phenomenon previously linked to aging. Furthermore, in the bone marrow of three-week-old offspring of high-fat diet-fed mothers, we identified a unique B cell population expressing CD19 and CD11b, and found increased expression of Cyclooxygenase-2 (COX-2) on myeloid CD11b, and on CD11b^hi^ B cells, with all the populations being significantly more abundant in offspring of dams fed HFD but not a regular diet. Altogether, we demonstrate that the offspring of obese mothers show metabolic and immune changes in the bone marrow at a very young age and prior to any symptomatic metabolic disease.

## INTRODUCTION

In many life-threatening diseases, such as cancer and cardiovascular ailments, patient outcomes can be improved by early detection. Obesity and metabolic diseases are the biggest epidemics in history, and not only increase mortality but also the risk of cancer and cardiovascular diseases, thus posing major challenges to healthcare systems worldwide (1). Moreover, obesity rates are rising, and disproportionally affect underserved and vulnerable racial, ethnic, and socioeconomic communities (2). While family history plays an important role, only 20% of obesity and metabolic disease cases are explained by genomic variations (3, 4).

Approximately 30% of US women are obese immediately prior to pregnancy (5). Epidemiological studies have revealed a strong association between obesity in pregnancy and adverse long-term consequences for offspring health and well-being (6); in particular, greater maternal pre-pregnancy body mass index (BMI) is linked to pregnancy complications (7) and progression of cardiovascular and metabolic diseases during the offspring’s adult life, including obesity, hyperinsulinemia, lipotoxicity, and increased metabolic inflammation (8–11). This intergenerational transmission may partly explain the present exacerbated epidemic of obesity and metabolic diseases (12). In a recently published study, prenatal metabolomic analysis of maternal plasma revealed several metabolites that might potentially mediate the programming effect of maternal obesity, including amino acids, sterols, and nucleotides (12). Thus, the individual’s risk of obesity and metabolic disease can be influenced during the earliest periods of their life, embryonic development, and recognizing this creates an opportunity for truly early detection of these disorders.

Accumulating epidemiological evidence and data from animal models clearly demonstrate that maternal obesity is accompanied by metabolic inflammation, an increase in adipose tissue macrophages, and systemic elevation of cytokine levels (13). Data from our lab and others demonstrate that this form of inflammation extends to the placenta, indicating that maternal obesity exposes the fetus to chronic inflammation during gestation (14–16). This might explain why neonates born to obese mothers are at increased risk of bacterial sepsis and necrotizing enterocolitis (17, 18). Moreover, it has been reported that immune system-related complications persist into adult life, placing the offspring of obese mothers at high risk of respiratory infections (19, 20), asthma (21), wheezing (22), type 2 diabetes (23), and even cancer (24).

The bone marrow (BM) is the primary site of hematopoiesis, and various physiological and pathological characteristics of the BM regulate the proliferation and migration of hematopoietic progenitors through blood into tissues (25). Published data reveal that the metabolic environment affects bone marrow activation, thereby leading to cardio-metabolic diseases and increased inflammation. Obesity, for example, has been linked to dysregulations in bone marrow homeostasis (26), and a mouse model of dyslipidemia and obesity reported elevated bone marrow hematopoietic activity and myeloid progenitor proliferation and expansion (27–29). Furthermore, diabetes mellitus in mice is also associated with expansion of BM myeloid progenitors (30, 31), myeloid cells and hematopoietic stem cells have been reported to undergo metabolic remodeling in adaptation to a changing metabolic environment (32–34). What effect maternal obesity has on bone marrow metabolism in offspring remains to be determined.

We have recently generated and reported a mouse model of maternal high-fat diet (HFD)-feeding that recapitulates the metabolic dysregulations seen in humans born to obese mothers, including increased adiposity, glucose intolerance, asthma, and insulin resistance (35, 36). Here we conducted targeted metabolomic profiling of bone marrow cells in our mouse model of obesity-induced developmental programming, comparing profiles of newly-weaned offspring from mothers fed a high-fat diet to those from mothers fed a normal diet to identify changes in offspring bone marrow that carry increased risk for developing obesity-related metabolic disorders later in life. We also analyzed the immune complexity of offspring bone marrow and blood tissues to identify a novel set of surrogate biomarkers that aid predicting risk of future obesity-related disorders.

## MATERIALS AND METHODS

### Study approval

All animal experiments were approved by the Oregon Health & Science University’s Institutional Animal Use Committee (Protocol # IP00432).

### Mouse model of maternal obesity

All studies were performed using wild-type FVB/N mice. Mice were kept under a 12-hour light/dark cycle, between 18-23 degrees Celsius with 40-60% humidity, in stress-free/bacteria-free conditions. Mice were caged in groups of three to five whenever possible. Food and water were given ad libitum. Body weights were collected weekly. In this investigation of diet-induced maternal obesity, a high-fat diet (Teklad Cat#TD.06415) or the control regular diet (RD, PicoLab® Laboratory Rodent Diet Cat # 5L0D) was given to virgin female FVB/NJ mice from six weeks of age and throughout the entire study. The composition of these diets has been reported previously (35, 36). After eight weeks of dietary intervention, RD-and HFD-fed female mice were bred to an age-matched RD-fed male. At weaning, male and female offspring from each mother were randomly selected for study. Every mouse in each experimental group belongs to a different litter. The offspring were fed the regular diet only, starting from weaning and throughout their life.

### Body fat mass quantitation using EchoMRI

Body composition was measured using an EchoMRI 4-in-1/1000 as we described previously(36).

### Preparation of bone marrow cells

Bone marrow cells were isolated as previously reported (37). Briefly, femurs were flushed using a 25G5/8 needle with medium containing RPMI1640 + 2% FBS + 10 units/ml heparin + penicillin and streptomycin. Bits of bones were removed using a sterile 4 mm nylon cell strainer (Falcon 352340). Cells were dissolved in 50 ml medium and centrifuged at 2000 rpm (900 x *g*) for 10 minutes at 4°C. Cell pellets were then washed twice with 50 ml of serum-free RPMI (RPMI1640 + 20 mM Hepes + penicillin and streptomycin, adjusted to pH 7.4 before filtering), centrifuged at 2000 rpm at 4°C for 5 minutes, and resuspended in 25 ml of serum-free RPMI.

### Metabolomics

Bone marrow cells collected from 3-week-old offspring of regular or high-fat diet-fed mothers were subjected to targeted metabolomics analysis. Cell metabolites were identified and analyzed by the West Coast Metabolomics Center at UC Davis. The metabolomics data are available at the NIH Common Fund’s National Metabolomics Data Repository (NMDR) website, the Metabolomics Workbench, https://www.metabolomicsworkbench.org, where they have been assigned Study ID ST002803.

### Sample processing for mass spectrometry

Each full sample was extracted using the Matyash extraction procedure which includes MTBE, MeOH, and H_2_O. The organic (upper) phase was dried down and submitted for resuspension and injection in liquid chromatography (LC) while the aqueous (bottom) phase was dried down and submitted to derivatization for gas chromatography (GC).

#### a. Gas chromatography – mass spectrometry

We used a 7890A GC coupled with a LECO TOF. About 0.5 mL of derivatized sample was injected using a splitless method into a RESTEK RTX-5SIL MS column with an Intergra-Guard at 275°C with a helium flow of 1.0 mL/minute. The GC oven was set to hold at 50°C for 1 minute, then ramp at 20°C/min to 330°C, and finally hold for 5 min. The transfer line was set to 28 °C, while the ion source was set to 250°C. Data were collected from 85 m/z to 500 m/z at an acquisition rate of 17 spectra/sec.

#### b. Liquid chromatography – mass spectrometry

Samples were resuspended with 110 mL of a solution of 9:1 methanol:toluene and 50 ng/mL CUDA, shaken for 20 seconds, sonicated for 5 minutes at room temperature, and then centrifuged for 2 minutes at 16100 rcf. Approximately 33 mL was aliquoted into in a vial with a 50 mL glass insert for positive mode metabolomics. The samples were then loaded up on an Agilent 1290 Infinity LC stack. The positive mode was run on an Agilent 6530 with a scan range of m/z 120-1200 Da and acquisition speed of 2 spectra/s. Between 0.5 and 2.0 mL was injected onto a Waters ACQUITY UPLC CSH C18 1.7 mm 2.1x100 mm column with a VanGuard PreColumn (2.1 x 5 mm). The gradient used was 0 minute 15% (B), 0–2 minute 30% (B), 2–2.5 mininute 48% (B), 2.5–11 minute 82% (B), 11–11.5 minute 99% (B), 11.5–12 minute 99% (B), 12–12.1 minute 15% (B), 12.1–15 minute 15% (B) with a flow rate of 0.6 mL/minute. The mass resolution for the Agilent 6530 is 10,000 for ESI (+).

### Measuring the mitochondrial function of bone marrow cells

#### Reagents

cRPMI: RPMI without phenol red (11835030, Gibco) and with 1% Pen/Strep (15140122, Gibco) and 10% FBS (35-010-CV, Corning). Seahorse calibrant (100840–000). Seahorse DMEM basal media (103575–100) was supplemented with 25 mM glucose, 1.0 mM pyruvate, and 4.0 mM glutamine; the pH was adjusted to 7.4 immediately prior to use.

#### Procedure

Bone marrow cells were harvested from the femur and tibia of euthanized mice as previously reported (37). Tissues were removed to expose bone and the ends of the bones were cut, after which the marrow was flushed with PBS using a 23-gauge needle and syringe; the resulting flushate was passed through a 70 mm filter. RBCs were lysed by incubating with RBC lysis buffer for 10 minutes at RT. The remaining non-RBC cells were washed, frozen, and stored at -80°C until the day of assay. The day before the assay, a Seahorse sensor plate (Agilent) was calibrated overnight in a non-CO_2_ incubator. On the day of the assay, frozen cells were thawed, counted according to staining with trypan blue, plated at 200,000 cells/well in a poly-D-lysine-coated plate, and let rest at 37°C for 1 hour in RPMI. The medium was then replaced with supplemented Seahorse basal medium, and plates calibrated in a non-CO_2_ incubator for 45 minutes prior to experiment. Drugs were added to cartridge plates at the following final concentrations: for the ATP assay, 2 mM oligomycin, 0.5 mM rotenone, and 0.5 mM antimycin A; and for the Mito Stress Assay, 2.0 mM oligomycin, 2.0 mM FCCP, 0.5 mM rotenone, and 0.5 mM antimycin A. ATP Assay and Mito Stress Assay programs were run on a SeahorseXFe96 (Agilent, Santa Clara, CA) as we previously described (38).

### Flow cytometry

The antibody list is presented in the Supplemental Materials (Table S1). In preparation for flow cytometry, nearly 10^6^ cells were first incubated on ice for 30 minutes in a solution consisting of a 1:10 Fc Receptor Binding Inhibitor (eBiosciences) and 1:500 Live/Dead Aqua stain (Invitrogen) diluted in PBS. Cells were then combined with fluorescently-labeled monoclonal antibodies as previously described (39) in a solution of PBS + 5% FCS + 1.0 mM EDTA (FACS buffer). After another 30 minutes incubation on ice, the cells were washed 1x with FACS buffer. Next, the stained cells were treated with permeabilization/fixation buffer (eBioscience) on ice for 10 minutes, then washed 1x using permeabilization buffer (eBioscience). Next, the permeabilized cells were subjected to intracellular staining: cells were incubated with fluorescently-labeled monoclonal antibodies for intracellular staining in a solution of PBS + 5% FCS + 1.0 mM EDTA (FACS buffer) for 30 minutes on ice. Flow cytometry data was acquired on a Cytek Aurora (Cytek Biosciences) spectral flow cytometer and the data analyzed using FlowJo software v9.5.

#### Quantitative PCR analysis

Total RNA was isolated using the RNeasy mini kit (Qiagen) with an additional DNase treatment using the DNA-free™ Kit (Thermo Fisher, Ref: AM1906); the yield was quantified with a Nanodrop (Thermo Fisher). cDNA was prepared from 100 ng of total RNA by reverse transcription PCR (RT–PCR) using a High-Capacity cDNA Reverse Transcription Kit (Thermo Fisher, Ref: 4368814) per the manufacturer’s instructions. Quantitative PCR to detect *IL-6*, *TNFa*, and *IFNγ* mRNA was performed on the StepOne Plus Real Time PCR System instrument using the QuantiTect SYBR Green PCR Master Mix (Qiagen, Cat No: 204143). The primers are presented in the Supplemental Materials (Table S2). Fold changes in expression were calculated by the 2^(-ΔΔCt)^ method, using ACTB as the endogenous control.

### Statistics

Statistical analysis was performed using Prism 9 software (GraphPad, San Diego, US). Data are expressed as mean ± SEM (standard error of the mean). The Shapiro–Wilk test was used to test for normal distribution in all data sets. Significant differences between the groups in normally distributed data were tested as an interaction between fetal sex (M, F) and maternal diet (regular vs. high fat) within a two-way ANOVA followed by unpaired two-tailed Student’s *t*-test. For non-parametric data, the Kruskal–Wallis test was applied followed by the Mann–Whitney U post hoc test. For the offspring, sample size represents the number of mothers fed either a regular or high-fat diet and is represented in graphs as one dot per sample. Flow cytometry and RT-PCR plots are representatives of at least three replicates.

#### Metabolomics data analysis

Metabolites were filtered to include only named ones present in 5% of samples, with a detection rate of >70% across 40 quality control (QC) pooled samples, relative standard deviation <15%, and a D-ratio of the standard deviation of QC samples divided by the standard deviation of samples <50%. Robust standardization (median centered and divided by the standard deviation) was performed on the log-transformed metabolite levels. Outliers greater than 5 standard deviations were clipped. Multiple comparisons were controlled using Benjamini-Hochberg’s false discovery rate. Comparison between the groups was conducted using two-way ANOVA to identify metabolites that differ as a result of maternal diet and fetal sex. Profiling was conducted using the MetaboAnalyst software. Principal component analysis (PCA) of metabolite profiles was utilized to analyze overall variance among mice. Pathway sets were obtained from databases such as the Kyoto Encyclopaedia of Genes and Genomes (KEGG), Reactome, or Ingenuity Pathway Analysis.

## RESULTS

### Mouse model of maternal obesity

Six-week-old females were placed either on a regular (13% kcal from fat) or a high fat (45% kcal from fat) diet (RD and HFD, respectively) for 8 weeks prior to pregnancy and throughout pregnancy and lactation (**Fig. 1A** and (40)). The composition of the diets were reported previously (35, 36) . After 8 weeks, the HFD-fed females evidenced no change in body weight when compared with RD-fed females (**Table 1**). However, body fat percentage as determined by echo magnetic resonance, was increased in HFD-fed females by 54% (*p*=0.03) (**Table 1**). After 8 weeks on diet, RD- and HFD-fed females were bred to RD-fed males. By term, the HFD-fed dams had a 1.5-fold increase in gestational weight gain in relation to each pup (*p*<0.1) and had 10% weight retention post-weaning (*p*=0.04, **Table 1**).

**Figure 1.**
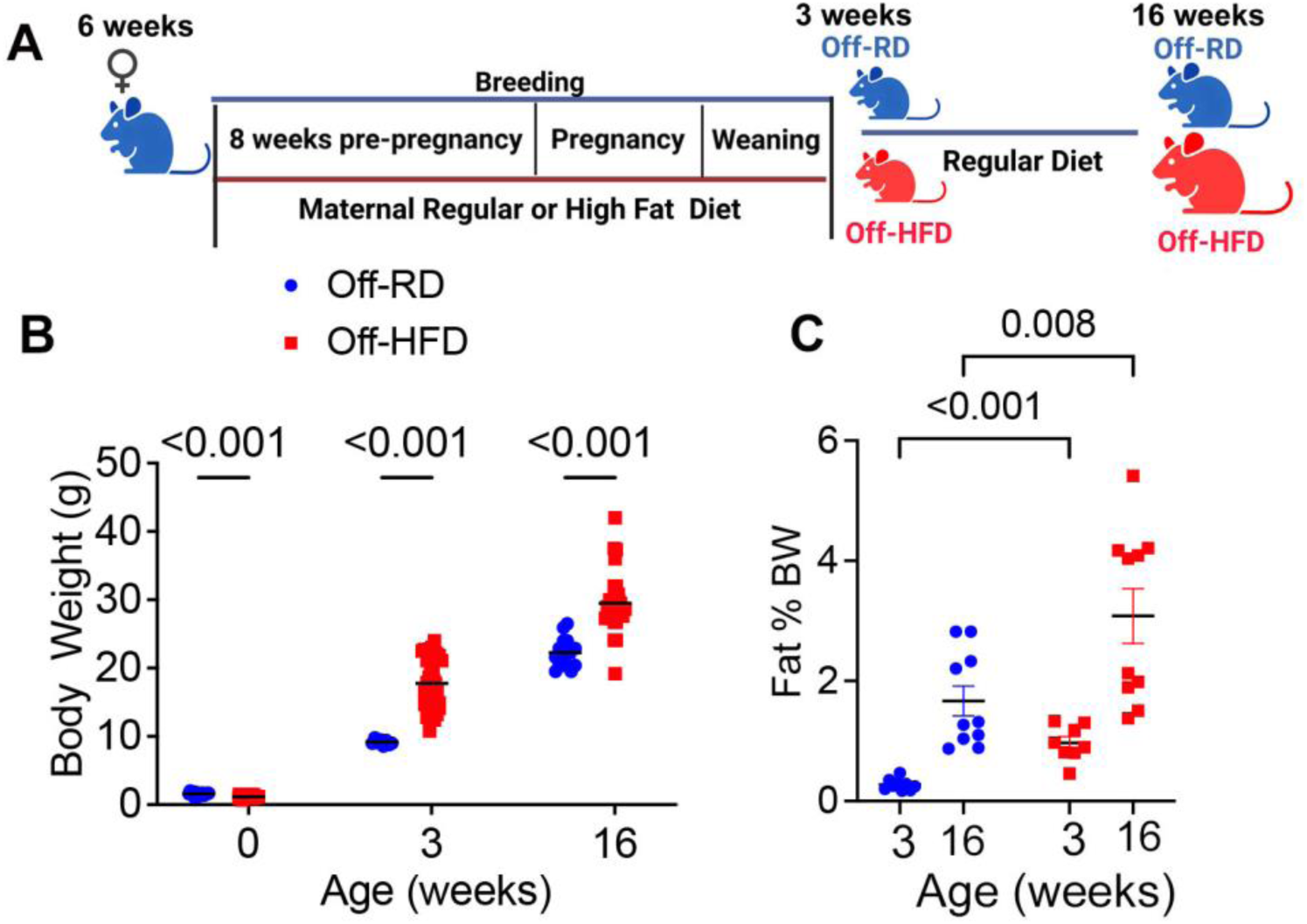
Mouse model of maternal diet-induced adiposity. **A.**, Experimental design. **B**, Body weights of offspring at birth (0), 3 weeks and 16 weeks of age. **C**, Body fat percentage in 3- and 16-week-old male and female offspring measured by ECHO MRI. Data are shown as mean ± standard error of the mean. N=8-35/sex/maternal diet intervention/age; *p*-values are shown on graphs. Cartoon was created with BioRender.com.

**Table 1.** Characteristics of regular diet (RD)- or high-fat diet (HFD)-fed mothers. BW, body weight; N=5-17/group of dietary intervention; Data presented as mean ± SEM.

| Characteristics | RD-fed | HFD-fed | *p*-value |
| --- | --- | --- | --- |
| Starting BW (g) | 20.81±0.67 | 21.52±0.18 | ns |
| Pre-pregnancy BW (g) | 24.23±0.82 | 24.14±0.46 | ns |
| Pre-pregnancy maternal adiposity (%) | 22.90±1.44 | 34.55±1.62 | 0.03 |
| Gestational weight gain/pup (g/pup) | 0.75±0.06 | 1.11±0.14 | 0.05 |
| Post-weaning BW (g) | 29.6±0.55 | 32.18±1.25* | 0.04 |

Consistent with our prior report (36), the offspring of mothers on a high-fat diet (Off-HFD) were born with 28% smaller birth weights (**Fig. 1B**); however, they evidenced catch-up growth during lactation, and by weaning (3 weeks) and at adult age (16 weeks) were significantly heavier compared with the offspring of dams on a regular diet (Off-RD, p<0.001 ). In addition, the adiposity of the Off-HFD was significantly increased at 3 and 16 weeks of ages vs. age-matched Off-RD mice (**Fig. 1C**, *p*<0.05).

### Maternal high-fat diet affects bone marrow metabolism

Dyslipidemia in humans leads to abnormal metabolism in bone marrow cells (41), and mouse models of metabolic diseases have likewise demonstrated changes in bone marrow activation (27–31). Our team has previously reported dyslipidemia in newborn babies born to obese mothers (11), and our mouse model presents progressive obesity and metabolic dysregulations in the offspring of HFD-fed mothers. Accordingly, we leveraged this model to investigate the effects of maternal obesity on offspring bone marrow cell metabolism by conducting a metabolomics analysis of bone marrow cells from 3-weeks-old, newly weaned offspring of dams fed regular and high-fat diets (**Fig. 2A**).

**Figure 2.**
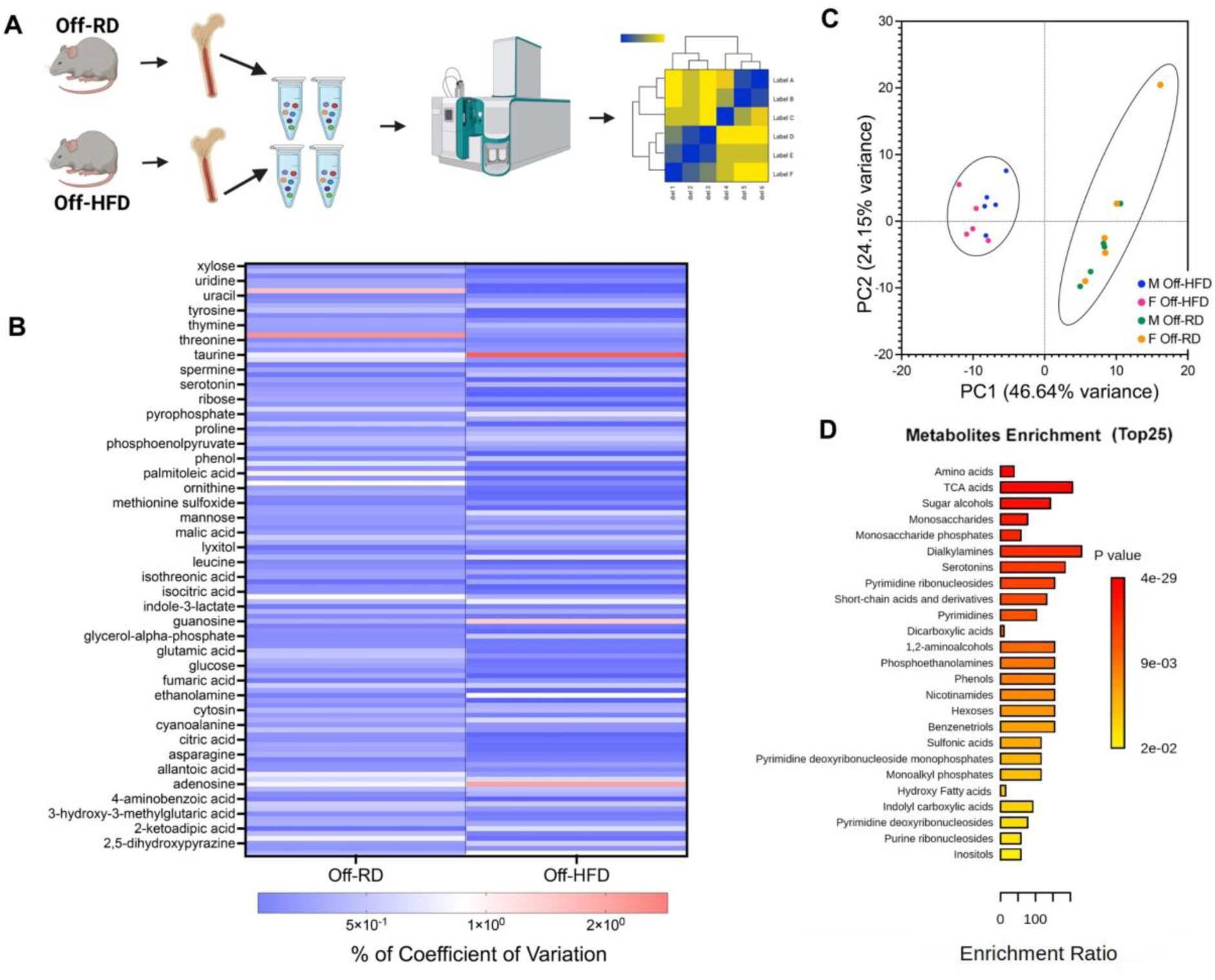
Metabolomic analysis of bone marrow cells. **A**, Experimental workflow. **B,** Heatmap of differentially expressed metabolites identified in bone marrow of 3-week-old Off-RD and Off-HFD mice. Male and female data are pooled. **C**, Principal component analysis (PCA) of the metabolomics dataset obtained from bone marrow cells showing discrimination of samples collected from male and female offspring of RD-(green and orange, respectively) and HFD-fed (blue and red, respectively) mothers. No sex-specific separation was detected. Significantly enriched metabolic pathways identified using the MetaboAnalyst 5.0 tool. The top 25 differentially expressed pathways are shown. N=5/group of maternal diet/sex; *p*<0.05. Cartoon was created with BioRender.com.

Overall, the analysis detected 122 annotated metabolites as significantly changed between the two groups (*p*<0.05, **Fig.2B**). Principal component analysis (PCA) of the metabolite abundances was able to discriminate between the two groups, with PCs 1 and 2 accounting for 46.64% and 24.16% of the variance (70.8% cumulative) between the groups (**Fig.2C**). Surprisingly, no sex-dependent differences were observed within the maternal diet groups.

In order to understand the coordinated changes among sets of metabolites, we performed metabolite set enrichment analysis, which revealed amino acid metabolism, the Krebs cycle, serotonin, and nitrogen metabolism to be among the metabolic pathways significantly impacted by maternal obesity (**Fig. 2D**).

### Maternal obesity affects bone marrow energy metabolism in the offspring of HFD-fed mothers

To expand upon the metabolomics data, we first analyzed changes in the central energy metabolism (glycolysis, Krebs cycle) of offspring bone marrow cells. Glycolysis starts with the uptake of glucose by transporters, and subsequent processing in the cytosol to yield pyruvate and lactate (**Fig. 3A**). We found the Off-HFD group to have significantly higher levels of the glycolysis intermediates glucose-6-phosphate (**Fig.3B**), fructose-6 phosphate (**Fig.3C**), 3-phosphoglycerate (**Fig. 3D**), phosphoenolpyruvate (**Fig. 3E**), and pyruvate (**Fig. 3G**), along with lower levels of lactate (**Fig. 3H**).

**Figure 3.**
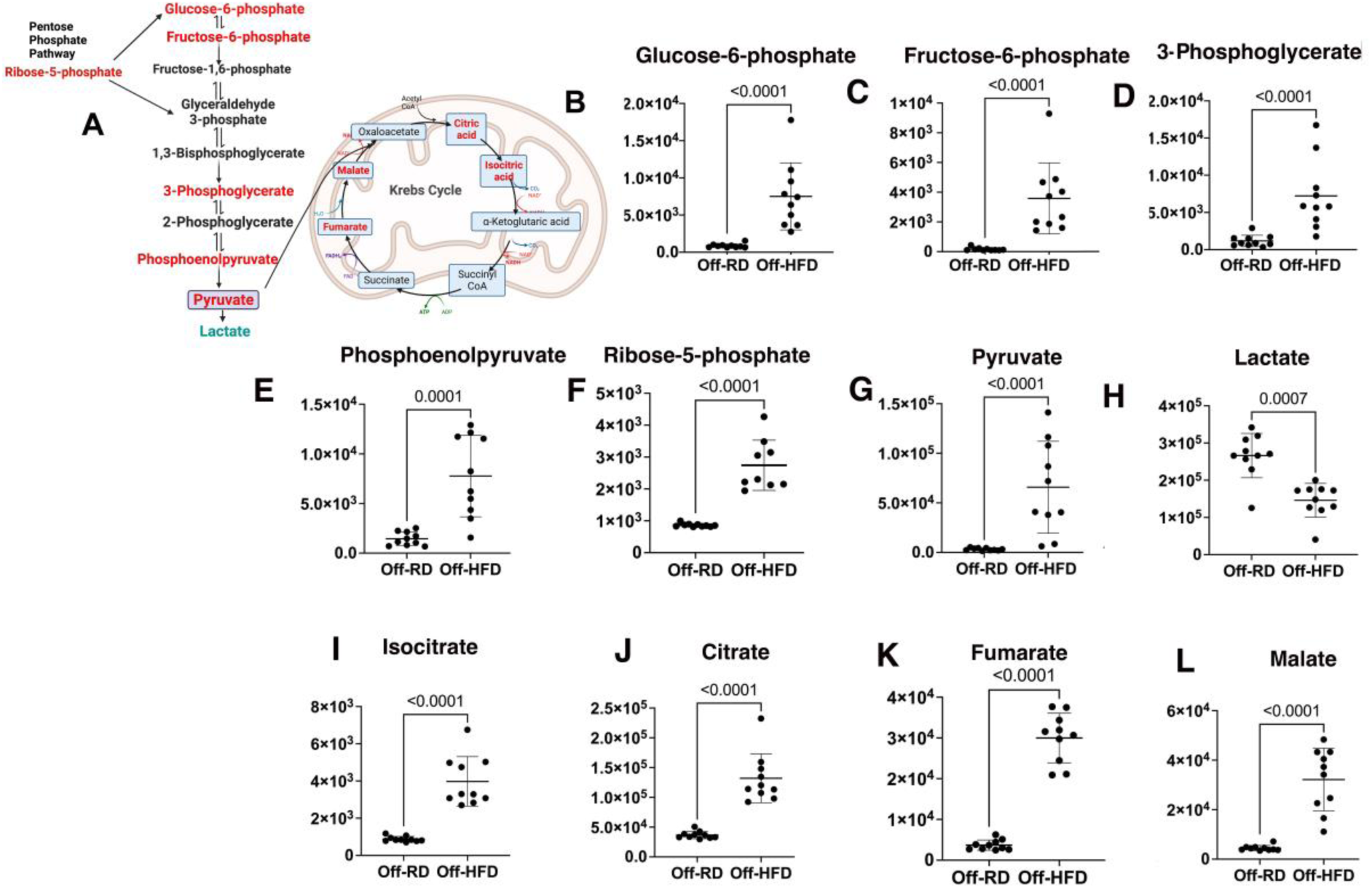
Changes in carbon metabolism and pentose phosphate pathway intermediates in the bone marrow of 3-weeks-old Off-HFD. **A**, Schematics of the pathways, red labeling indicates increase and green labeling – decrease in the intermediate levels. **B-H**, Glucose utilization is increased in the bone marrow of Off-HFD vs. off-RD mice, as shown by higher levels of glucose-6-phosphate (**B**), fructose-6-phosphate (**C**), and downstream metabolites 3-phosphoglycerate (**D**), phosphoenolpyruvate (**E**), ribose-5-phosphate (**F**), and pyruvate (**G**), and by lower levels of lactate (**H**). **I-L,** Increased levels of the Krebs cycle metabolites isocitrate (**I**), citrate (**J**), fumarate (**K**), and malate (**L**) in Off-HFD vs. Off-RD mice. Data are represented as mean ± SEM. N=5/group of maternal diet/sex; data from males and females were combined; *p*-values are shown on graphs. Cartoon was created with BioRender.com.

The pentose phosphate pathway (PPP) branches off from glycolysis and is a glucose-oxidizing pathway that plays a critical role in the homeostasis of vital organs and regulation of immune cell fate and functions ((42), **Fig. 3A**). Neutrophils and inflammatory macrophages use the oxidative branch of the PPP to generate reactive oxygen species (ROS) (43), and the non-oxidative branch for nucleotide biosynthesis and for linkage with glycolysis (44). Our metabolomics analysis revealed the Off-HFD to have a significant increase in ribose-5-phosphate, an intermediate in the non-oxidative PPP (**Fig. 3F**), and no evident change in the oxidative branch. These results indicated greater glucose utilization in Off-HFD bone marrow via activation of glycolysis and the PPP. Since the Off-HFD also exhibited lower levels of lactate, we predicted activation of the Krebs cycle and oxidative phosphorylation (OXPHOS). Indeed, several Krebs cycle intermediates were also affected by maternal obesity, with increased levels of isocitrate (**Fig. 3I**), citric acid (**Fig. 3J**), fumarate (**Fig. 3K**), and malate (**Fig. 3L**).

To assess the effect of maternal obesity on bone marrow oxidative phosphorylation, we measured bioenergetic profiles of bone marrow cells from Off-RD and Off-HFD mice using a Seahorse Analyzer (**Fig. 4A-B**). We found overall metabolic activity to be increased in the bone marrow of Off-HFD mice, evidenced by enhanced production of ATP (**Fig. 4C**), and a predominant increase in OXPHOS over glycolysis (**Fig. 4D-E**). Further examination of OXPHOS was conducted using the mitochondrial inhibitors oligomycin (ATP synthase inhibitor), FCCP, and rotenone with antimycin A (electron transport complex I and III inhibitors). This revealed that basal, ATP-induced, and maximal respiration, as well as spare capacity, were all increased in the bone marrow of 3-weeks-old Off-HFD mice compared to Off-RD mice (**Fig. 4F-I**).

**Figure 4.**
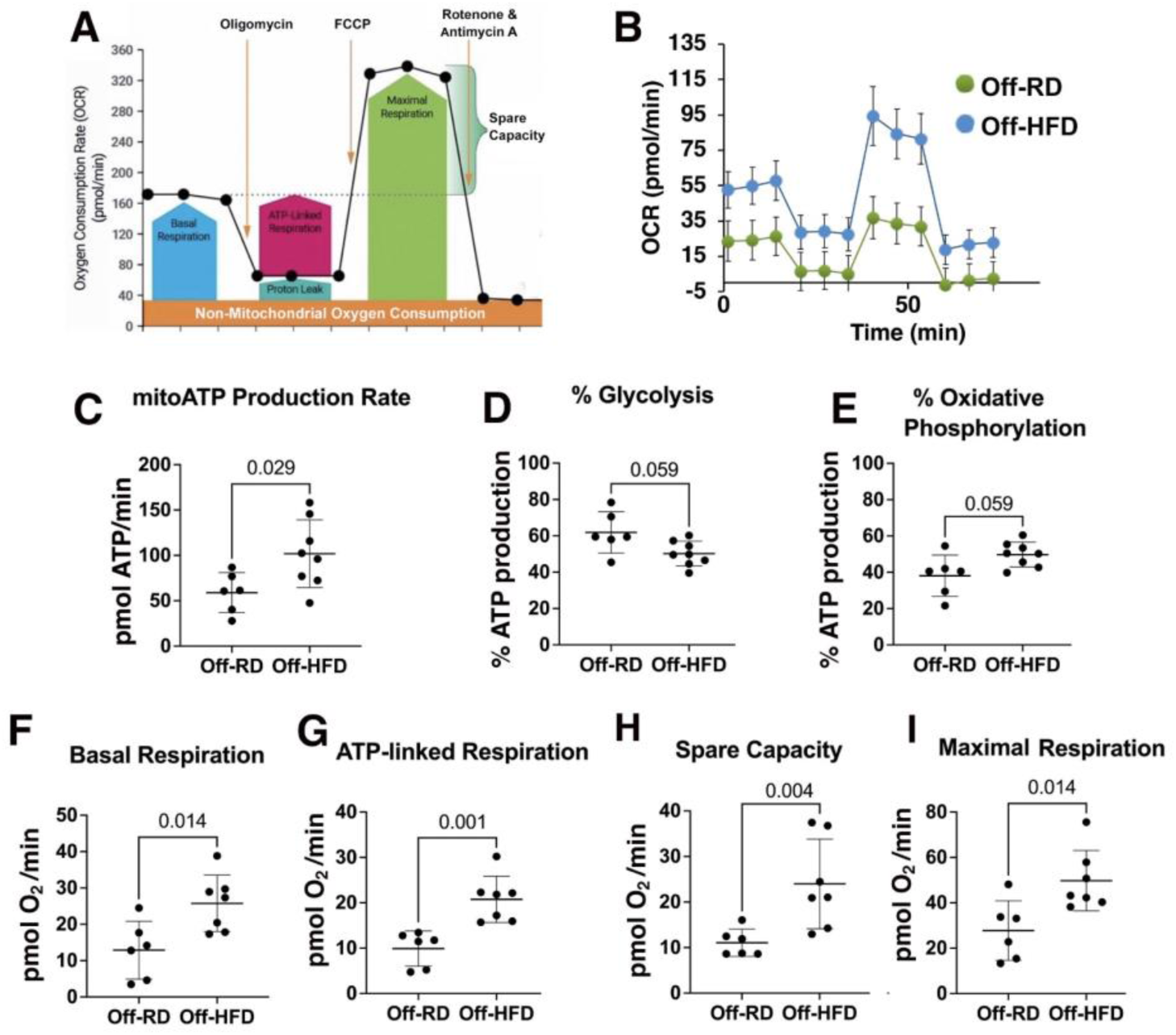
Increased mitochondrial respiration in the bone marrow cells of 3-weeks-old Off-HFD. **A-B**, Oxygen consumption rate (OCR) was measured in cultured bone marrow cells from 3-weeks-old Off-RD and Off-HFD using the Seahorse XF Cell Mito Stress Test. Measurements were taken before and after treatment with oligomycin (1.0 µmol/L), FCCP (1.1 µmol/L), and finally antimycin A/rotenone (0.5 µmol/L each), and the data were analyzed with the Wave 2.6 software. Overall workflow and representative graphs are shown in (**A**) and (**B**), respectively. **C**, Rate of ATP production from oxidative phosphorylation (OXPHOS). **D-E**, Percent ATP produced through glycolysis (**D**) and OXPHOS (**E**). **F**, Basal respiration; **G**, ATP-linked respiration; **H**, Spare respiratory capacity; and **I**, Maximal respiration. Data are represented as mean ± SEM. Data from males and females were pooled; N=3/sex/maternal diet; *p*-values are shown on the graphs.

### Maternal obesity affects bone marrow amino acid metabolism in the offspring of HFD-fed mothers

In addition to alterations in energy cycle, the metabolomics data revealed changes in levels and metabolism of important amino acids. In general, we found that several amino acids, except glutamate and glycine, were reduced in the Off-HFD group relative to the Off-RD one (**Table 2**).

**Table 2.**
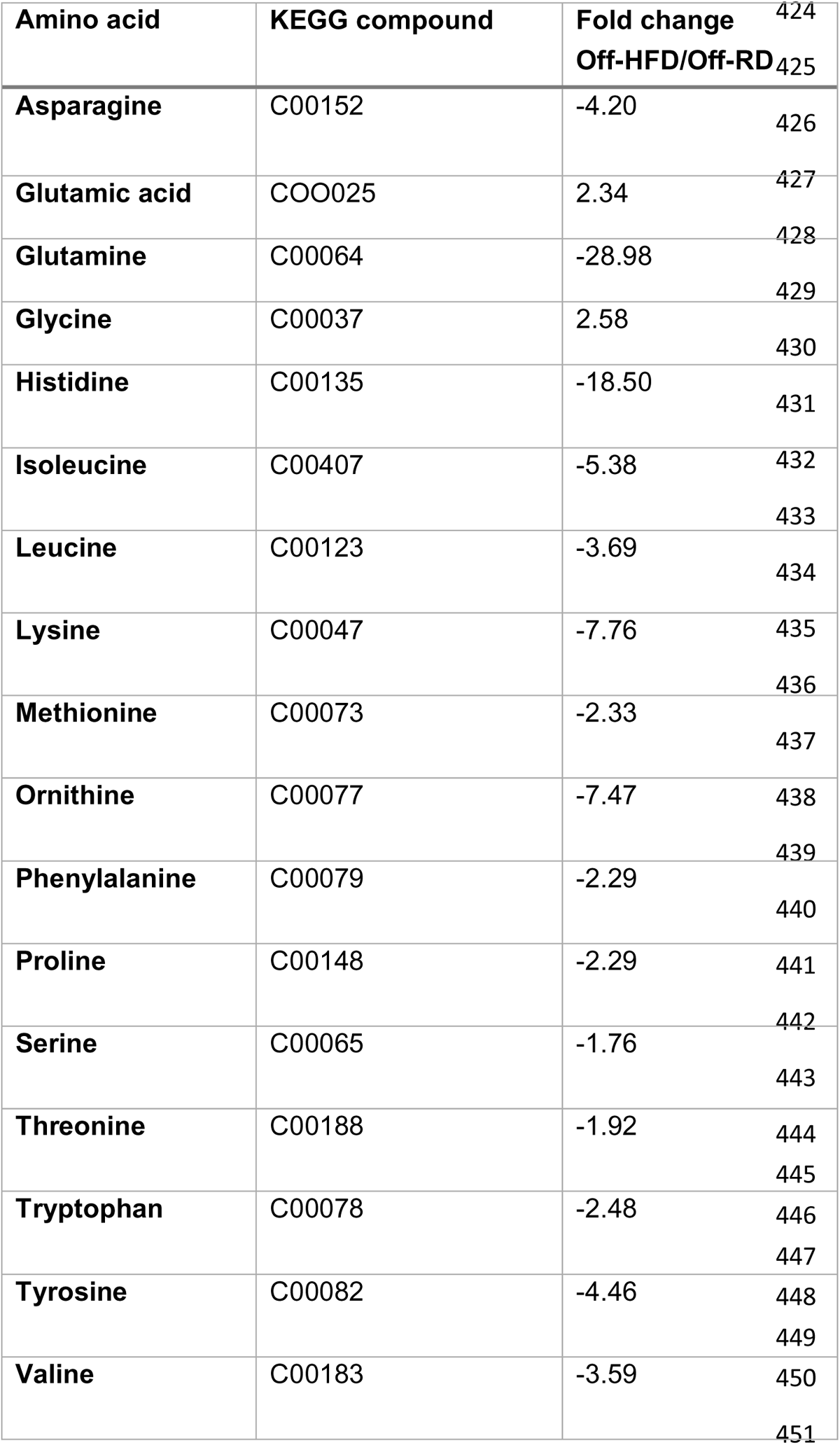
List of essential and non-essential amino acids significantly changed in the bone marrow cells of 3 weeks-old Off-HFD vs. Off-RD. Data expressed as fold change in Off-HFD relative to sex-matched Off-RD. N=5/sex/per group; *p*<0.05 vs. Off-RD.

Tryptophan, an essential amino acid, and its downstream metabolites were previously linked to modulation of immune function, control of hyperinflammation, and induction of long-term immune tolerance (45, 46). Cancer, chronic inflammatory, and autoimmune diseases are associated with a decline in tryptophan metabolism during aging and age-related diseases. Serotonin, a product of tryptophan metabolism (**Fig.5A**), is known for its role as a neurotransmitter (47). However, peripheral serotonin has been found to play an important role in inflammation and immunity (48), specifically in regulation of T cells, macrophages, dendritic cells, and platelets, and release of inflammatory cytokines. Our data identified tryptophan levels in the bone marrow of 3-weeks-old Off-HFD to be significantly decreased (**Fig. 5B**), while levels of serotonin (**Fig. 5C**) increased vs. Off-RD mice (*p*<0.05).

**Figure 5.**
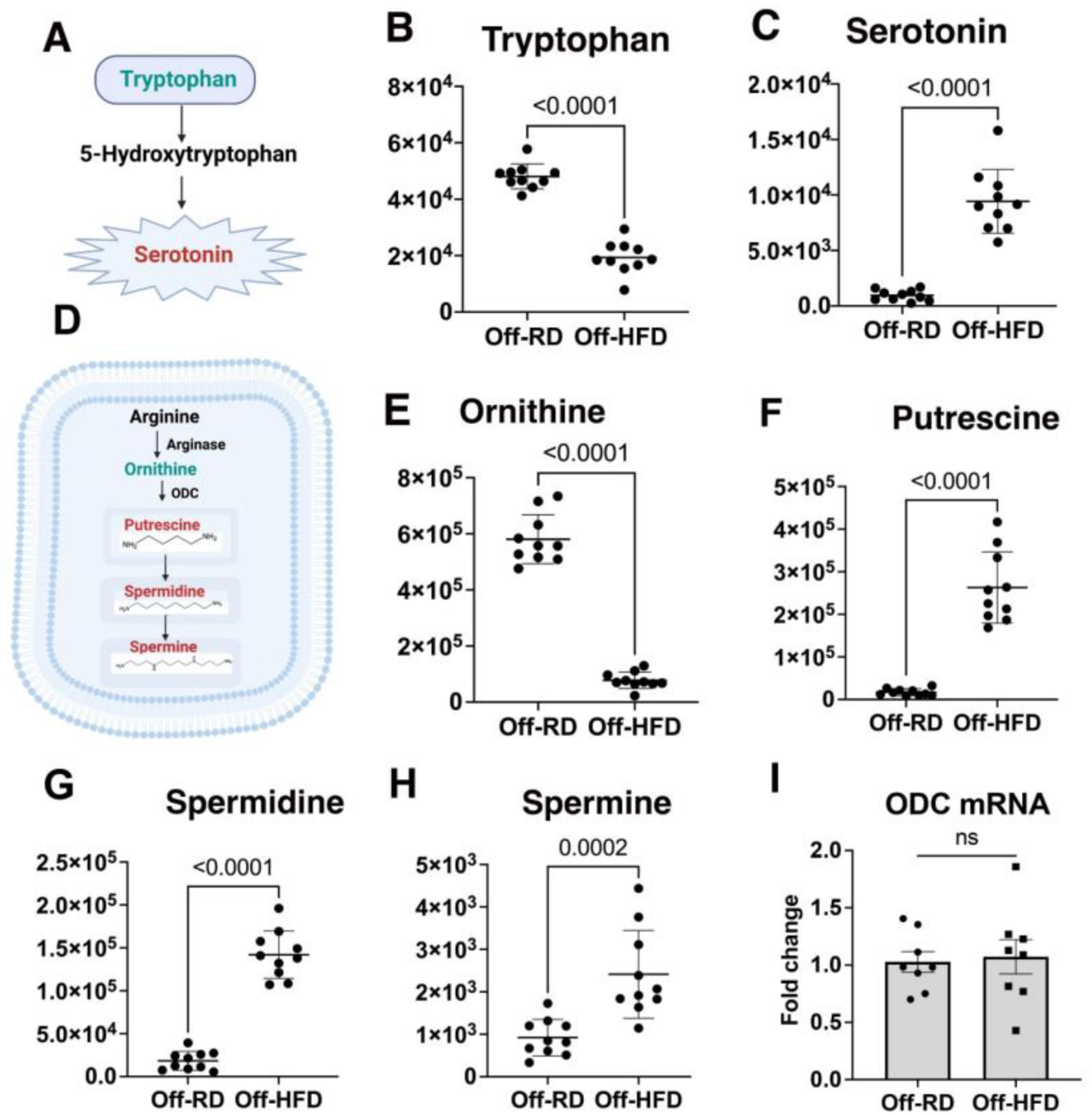
Dysregulation of tryptophan and arginine metabolism in the bone marrow of 3-weeks-old Off-HFD. **A**, Schematic representation of tryptophan metabolism. **B-C,** Measured levels of tryptophan (**B**), and serotonin (**C**). **D-I**, Shunting of arginine metabolism into polyamine synthesis. **D,** Schematic representation of arginine metabolism. **E**, Decreased levels of ornithine in Off-HFD vs. Off-RD mice. **F-H**, Increased levels of the polyamines putrescine (**F**), spermidine (**G**), and spermine (**H**) in Off-HFD vs. Off-RD mice. **I**, Quantification of ornithine decarboxylase (*ODC*) mRNA using RT-PCR. Data are represented as mean values ± SEM. Data from males and females were combined; N=5/ sex/maternal diet; *p*-values are shown on the graphs. Cartoons were created with BioRender.com.

Arginine is catabolized by five different groups of enzymes, including arginase II, for the synthesis of ornithine, proline, and glutamate (49), and approximately 2% of metabolized arginine is utilized for polyamine synthesis (**Fig.5D**). Published data suggest that polyamines play a critical role in modulation of immune cell function (50). Specifically, polyamines inhibit lymphocyte proliferation, reduce neutrophil locomotion, and suppress macrophage-mediated tumoricidal activity via reprogramming proinflammatory to anti-inflammatory phenotypes (51). In addition, blood polyamine levels are reported to be increased in childhood obesity, indicating a role for polyamine metabolism in the settings of obesity (52). Arginine concentrations in the bone marrow of 3-weeks-old Off-RD and Off-HFD mice exhibited no differences (not shown), but downstream ornithine (**Fig. 5E**) levels were decreased significantly in the Off-HFD (p<0.0001). Ornithine-derived polyamines—putrescine, spermidine, and spermine—were all significantly increased in Off-HFD vs. Off-RD mice (*p*<0.005, **Fig. 5F-H**). Since biosynthesis of ornithine-derived polyamines is regulated by the enzyme ornithine decarboxylase (ODC), we quantified *ODC* mRNA by RT-PCR. Surprisingly, there was no significant difference between Off-RD and Off-HFD, which may indicate that the rise in polyamines is not caused by their increased production from ornithine (**Fig. 5I**).

### Maternal obesity affects immune complexity in the bone marrow of the offspring of HFD-fed mothers

We next utilized state-of-the-art spectral flow cytometry enabling highly multiplexed profiling of cell type identifiers to reveal diverse subtypes of immune cells. We first assessed immune complexity of the bone marrow from 3-weeks-old Off-HFD and Off-RD mice (**Fig.6A-B**) and found that maternal obesity did not affect overall immune complexity of offspring, albeit there was a significant increase in abundance of CD19^+^ B cells in Off-HFD (**Fig. 6C**). We further zoomed into the B cell population, which was increased in Off-HFD by utilizing the strength of our highly multiplexed analysis. We found CD11b^HI^B cells to be significantly increased in bone marrow of the Off-HFD, largely accounting for the observed increase in B cells (**Fig. 6D**). We also observed a significant increase in expression of cyclooxygenase-2 (COX-2) in myeloid CD11b^hi^ cells (**Fig. 6E**), and CD11b^hi^ B cells (**Fig.6F**). We further quired whether any of these changes in bone marrow output were also reflected in blood where we found a corresponding increase in CD19^+^ B cells as well as in CD11b^hi^B cells in Off-HFD vs. Off-RD mice (**Fig. 6G-H**). This finding is important as combinatorial analysis of blood can pave the way for future early detection of obesity-related metabolic disorders.

**Figure 6.**
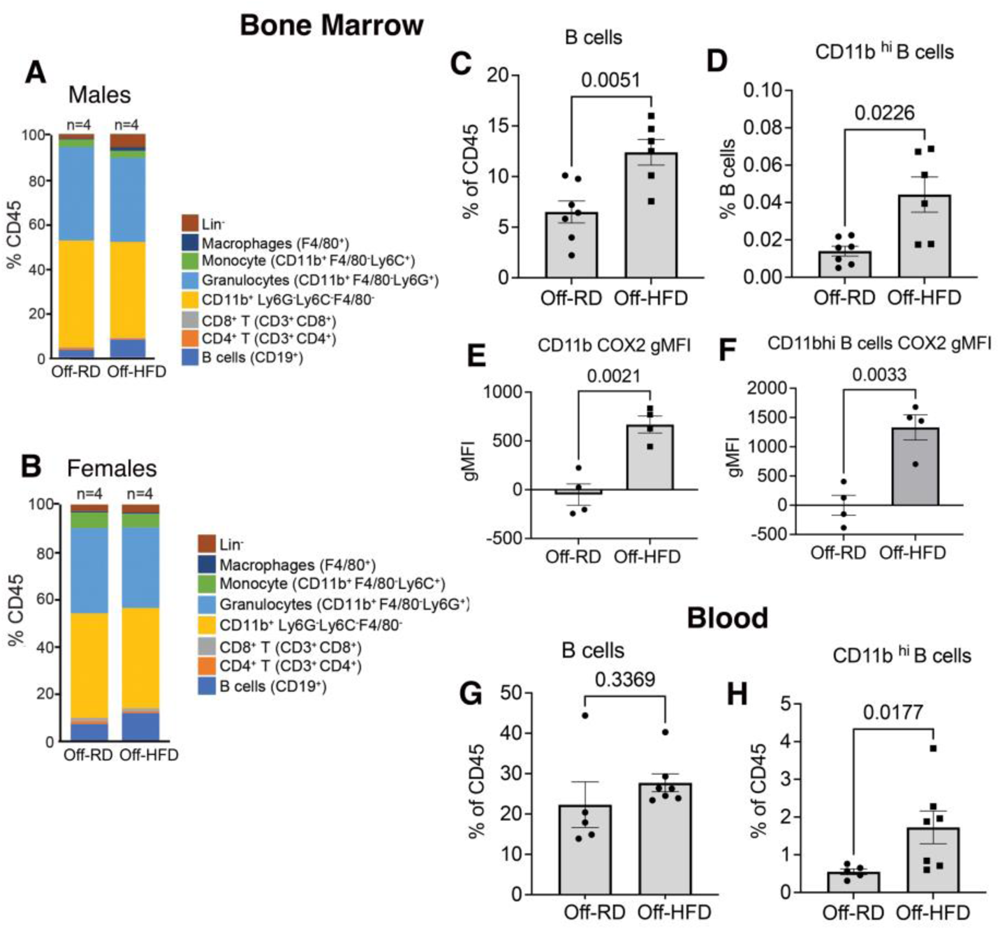
Increased B cell population and identification of a distinct subset in the bone marrow and blood of 3-weeks-old Off-HFD. **A-B**, Flow cytometric analysis of immune cell populations in the bone marrow of male (**A**) and female (**B**) 3 weeks-old offspring of RD-and HFD-fed mothers. Results are shown as the percentage of total CD45^+^ cells. **C-D,** Increase of B cell (**C**) and specifically CD11b^hi^ B cell (**D**) populations in the bone marrow of Off-HFD vs. Off-RD mice. **E-F**, Increase in COX2 expression on CD11b^+^ (**E**) and CD1^hi^ B-cells (**F**) cells. **G-H**, Increase of B cell (**G**) and CD11b^hi^ B cell (**H**) populations in the blood of 3-week old Off-HFD vs. Off-RD mice. Data are represented as mean values ± SEM. Data from males and females were combined; N=4-5/sex/maternal diet; *p*-values are shown.

Finally, we analyzed the gene expression of cytokines TNFa, IL-6, and IFNγ (**Supplemental Fig. 1**). This revealed expression of *TNFα* to be lower in Off-HFD than in Off-RD (*p*=0.028, **Suppl Fig. 1A**), whereas *IL-6* showed a tendency to increase that did not reach statistical significance (*p*=0.072, **Suppl. Fig. 1B**). No between-group change was observed in the expression of *IFN*γ mRNA (**Suppl. Fig. 1C**).

## Discussion

Obesity has become a significant socioeconomic burden worldwide (53, 54). While diet-inflicted obesity can be controlled by changing lifestyle, management of obesity due to genetic predisposition remains a challenge. In addition to genetic predisposition, maternal reprogramming of an embryo can also contribute to obesity and associated metabolic disorders (55–57). Overall, the immune system, specifically innate immune cells known to undergo functional state changes in tissue when exposed to various pro-inflammatory conditions, is increasingly recognized as rate-limiting in the progression of metabolic disorders, including obesity (34, 58). Moreover, bone marrow cells were recently found to be involved in response to nutritional stress (59). In this study, we sought to analyze early life changes in the hematopoietic system of offspring of obese mouse dams with the hope of identifying key targets for potential therapeutic intervention or diagnostic biomarkers for early detection of at-risk populations. We specifically examined the hematopoietic system and circulating blood of 3-weeks-old newly weaned mice to identify changes relative to offspring of normal dams (Off-RD), and performed a multimodal analysis to allow corroboration of data across different datasets. For instance, metabolomics data indicated increased Krebs cycle output in bone marrow cells of the Off-HFD, which was substantiated by measurements of oxidative phosphorylation taken with a Seahorse analyzer, which revealed increased O_2_ consumption. Activation of oxidative phosphorylation has previously been observed in aged bone marrow (60). Mitochondria are both a source and target of reactive oxygen species (61), and increased oxidative phosphorylation could potentially be a compensatory response to oxidative stress, a well-known feature of aging in other tissues (62). Our finding of reduced amino acid levels is also in agreement with aging bone marrow (60), aging rat brain tissue (63), and aging mammalian serum (64). A diminished amino acid pool in offspring of obese mothers could result from either reduced synthesis of amino acids or dysfunction of autophagy that would otherwise replenish the pool via protein turnover (65). Accordingly, our team has previously reported dysregulations in autophagy associated with intrauterine exposure to maternal obesity (66).

We also applied multiplexed flow cytometry to analyze immune complexity and phenotypes of bone marrow cells and revealed significant changes in immune composition. We identified a unique B cell type in the Off-HFD mice, characterized by increased expression of both CD19 and CD11b in bone marrow and blood. In autoimmune hepatitis, this B cell subtype has been noted to augment cytotoxic T cell responses, and *in vitro* studies revealed that the suppressive function of CD11b was mediated by impairment of T cell antigen receptor (TCR) signaling transduction and promotion of TCR downregulation (67). CD11b^hi^ B cells have been reported to be markedly elevated in lupus patients (68). however, in contrast to autoimmune hepatitis, CD11b^+^ B cells in lupus patients express more CD86 and have increased T cell– stimulating activity (75). Whether this B cell population is functionally relevant in the development of obesity and its associated metabolic disorders requires further careful investigation. In the short term, it will be interesting to determine if this change persists into adulthood, especially when obesity and disease phenotypes set in.

Our data also reveal an increase in COX2 expression on bone marrow myeloid cells of 3-week-old Off-RD. COX-2 is integral to macrophage phenotype and function (69); inhibition of COX-2 in cultured murine bone marrow cells results in enhanced differentiation to an antigen-presenting phenotype (70), and inhibition of human monocytes toward M2-type transcriptional skewing (71). In addition, COX-2 has been found to be highly expressed in chronic inflammatory disorders such as atherosclerosis (72), and in cancer, paracrine influences of tumor cell COX-2 potentiates mammary tumorigenesis (73, 74). The role of COX2 in the settings of maternal obesity remains unclear, and is a subject of ongoing investigation by our team.

Ultimately, our comprehensive analyses of bone marrow and blood in young animals using our model of maternal obesity confirm that systemic differences are established very early on in life. This understanding provides us with a unique opportunity to not only identify measures for early detection of this menacing disease, but also to develop novel preventive measures that can help millions of lives.

## Author Contributions

EP, YA, NG and SK – conducted all the experiments and analyzed the data; LK provided access to the Seahorse instrument and helped with Seahorse data analysis; SK – conducted flow cytometry experiments, and analyzed the data, AM-funding acquisition; SK, LMC and AM – conceptualization and design of the study, SK and AM wrote the manuscript. All authors provided feedback and assisted with preparing the final manuscript.

## REFERENCES

1. Moholdt T, Hawley JA. Maternal Lifestyle Interventions: Targeting Preconception Health. Trends Endocrinol Metab. 2020;31(8):561–9.

2. Bulik CM, Hardaway JA. Turning the tide on obesity? Science. 2023;381(6657):463-.

3. Mahajan A, Taliun D, Thurner M, Robertson NR, Torres JM, Rayner NW, et al. Fine-mapping type 2 diabetes loci to single-variant resolution using high-density imputation and islet-specific epigenome maps. Nat Genet. 2018;50(11):1505–13.

4. Locke AE, Kahali B, Berndt SI, Justice AE, Pers TH, Day FR, et al. Genetic studies of body mass index yield new insights for obesity biology. Nature. 2015;518(7538):197–206.

5. Driscoll AK, Gregory ECW. Increases in Prepregnancy Obesity: United States, 2016-2019. NCHS Data Brief. 2020(392):1–8.

6. Langley-Evans SC. Early life programming of health and disease: The long-term consequences of obesity in pregnancy. J Hum Nutr Diet. 2022.

7. Leddy MA, Power ML, Schulkin J. The impact of maternal obesity on maternal and fetal health. Rev Obstet Gynecol. 2008;1(4):170–8.

8. Schmatz M, Madan J, Marino T, Davis J. Maternal obesity: the interplay between inflammation, mother and fetus. Journal of Perinatology. 2010;30(7):441–6.

9. Segovia SA, Vickers MH, Gray C, Reynolds CM. Maternal obesity, inflammation, and developmental programming. Biomed Res Int. 2014;2014:418975.

10. Gutvirtz G, Wainstock T, Landau D, Sheiner E. Maternal Obesity and Offspring Long-Term Infectious Morbidity. J Clin Med. 2019;8(9).

11. Bucher M, Montaniel KRC, Myatt L, Weintraub S, Tavori H, Maloyan A. Dyslipidemia, insulin resistance, and impairment of placental metabolism in the offspring of obese mothers. J Dev Orig Health Dis. 2021;12(5):738–47.

12. Hu Z, Han L, Liu J, Fowke JH, Han JC, Kakhniashvili D, et al. Prenatal Metabolomic Profiles Mediate the Effect of Maternal Obesity On Early Childhood Growth Trajectories and Obesity Risk: the CANDLE Study. Am J Clin Nutr. 2022.

13. Basu S, Haghiac M, Surace P, Challier JC, Guerre-Millo M, Singh K, et al. Pregravid obesity associates with increased maternal endotoxemia and metabolic inflammation. Obesity (Silver Spring). 2011;19(3):476–82.

14. Challier JC, Basu S, Bintein T, Minium J, Hotmire K, Catalano PM, et al. Obesity in pregnancy stimulates macrophage accumulation and inflammation in the placenta. Placenta. 2008;29(3):274–81.

15. Muralimanoharan S, Guo C, Myatt L, Maloyan A. Sexual dimorphism in miR-210 expression and mitochondrial dysfunction in the placenta with maternal obesity. Int J Obes (Lond). 2015;39(8):1274–81.

16. Myatt L, Maloyan A. Obesity and Placental Function. Semin Reprod Med. 2016;34(1):42–9.

17. Suk D, Kwak T, Khawar N, Vanhorn S, Salafia CM, Gudavalli MB, et al. Increasing maternal body mass index during pregnancy increases neonatal intensive care unit admission in near and full-term infants. J Matern Fetal Neonatal Med. 2016;29(20):3249–53.

18. Rastogi S, Rojas M, Rastogi D, Haberman S. Neonatal morbidities among full-term infants born to obese mothers. J Matern Fetal Neonatal Med. 2015;28(7):829–35.

19. Griffiths PS, Walton C, Samsell L, Perez MK, Piedimonte G. Maternal high-fat hypercaloric diet during pregnancy results in persistent metabolic and respiratory abnormalities in offspring. Pediatric research. 2016;79(2):278–86.

20. Haberg SE, Stigum H, London SJ, Nystad W, Nafstad P. Maternal obesity in pregnancy and respiratory health in early childhood. Paediatr Perinat Epidemiol. 2009;23(4):352–62.

21. Dumas O, Varraso R, Gillman MW, Field AE, Camargo CA, Jr. Longitudinal study of maternal body mass index, gestational weight gain, and offspring asthma. Allergy. 2016;71(9):1295–304.

22. Guerra S, Sartini C, Mendez M, Morales E, Guxens M, Basterrechea M, et al. Maternal prepregnancy obesity is an independent risk factor for frequent wheezing in infants by age 14 months. Paediatr Perinat Epidemiol. 2013;27(1):100–8.

23. Morgan AR, Thompson JM, Murphy R, Black PN, Lam WJ, Ferguson LR, et al. Obesity and diabetes genes are associated with being born small for gestational age: results from the Auckland Birthweight Collaborative study. BMC Med Genet. 2010;11:125.

24. Eriksson JG, Sandboge S, Salonen MK, Kajantie E, Osmond C. Long-term consequences of maternal overweight in pregnancy on offspring later health: findings from the Helsinki Birth Cohort Study. Ann Med. 2014;46(6):434–8.

25. Lucas D. Structural organization of the bone marrow and its role in hematopoiesis. Curr Opin Hematol. 2021;28(1):36–42.

26. Benova A, Tencerova M. Obesity-Induced Changes in Bone Marrow Homeostasis. Frontiers in Endocrinology. 2020;11.

27. Nagareddy PR, Kraakman M, Masters SL, Stirzaker RA, Gorman DJ, Grant RW, et al. Adipose tissue macrophages promote myelopoiesis and monocytosis in obesity. Cell Metab. 2014;19(5):821–35.

28. Singer K, DelProposto J, Morris DL, Zamarron B, Mergian T, Maley N, et al. Diet-induced obesity promotes myelopoiesis in hematopoietic stem cells. Mol Metab. 2014;3(6):664–75.

29. Tolani S, Pagler TA, Murphy AJ, Bochem AE, Abramowicz S, Welch C, et al. Hypercholesterolemia and reduced HDL-C promote hematopoietic stem cell proliferation and monocytosis: studies in mice and FH children. Atherosclerosis. 2013;229(1):79–85.

30. Hoyer FF, Zhang X, Coppin E, Vasamsetti SB, Modugu G, Schloss MJ, et al. Bone Marrow Endothelial Cells Regulate Myelopoiesis in Diabetes Mellitus. Circulation. 2020;142(3):244–58.

31. Nagareddy PR, Murphy AJ, Stirzaker RA, Hu Y, Yu S, Miller RG, et al. Hyperglycemia promotes myelopoiesis and impairs the resolution of atherosclerosis. Cell Metab. 2013;17(5):695–708.

32. Mitroulis I, Ruppova K, Wang B, Chen LS, Grzybek M, Grinenko T, et al. Modulation of Myelopoiesis Progenitors Is an Integral Component of Trained Immunity. Cell. 2018;172(1-2):147–61.e12.

33. Mulder WJM, Ochando J, Joosten LAB, Fayad ZA, Netea MG. Therapeutic targeting of trained immunity. Nat Rev Drug Discov. 2019;18(7):553–66.

34. Netea MG, Joosten LA, Latz E, Mills KH, Natoli G, Stunnenberg HG, et al. Trained immunity: A program of innate immune memory in health and disease. Science. 2016;352(6284):aaf1098.

35. Calco GN, Alharithi YJ, Williams KR, Jacoby DB, Fryer AD, Maloyan A, et al. Maternal high-fat diet increases airway sensory innervation and reflex bronchoconstriction in adult offspring. Am J Physiol Lung Cell Mol Physiol. 2023;325(1):L66–l73.

36. Montaniel KRC, Bucher M, Phillips EA, Li C, Sullivan EL, Kievit P, et al. Dipeptidyl peptidase IV inhibition delays developmental programming of obesity and metabolic disease in male offspring of obese mothers. J Dev Orig Health Dis. 2022;13(6):727–40.

37. Liu X, Quan N. Immune Cell Isolation from Mouse Femur Bone Marrow. Bio Protoc. 2015;5(20).

38. Phillips EA, Hendricks N, Bucher M, Maloyan A. Vitamin D Supplementation Improves Mitochondrial Function and Reduces Inflammation in Placentae of Obese Women. Front Endocrinol (Lausanne). 2022;13:893848.

39. Ruffell B, Au A, Rugo HS, Esserman LJ, Hwang ES, Coussens LM. Leukocyte composition of human breast cancer. Proc Natl Acad Sci U S A. 2012;109(8):2796–801.

40. Montaniel KRC, Bucher M, Phillips EA, Li C, Sullivan EL, Kievit P, et al. Dipeptidyl peptidase IV inhibition delays developmental programming of obesity and metabolic disease in male offspring of obese mothers. J Dev Orig Health Dis. 2022:1–14.

41. Tall AR, Yvan-Charvet L, Westerterp M, Murphy AJ. Cholesterol efflux: a novel regulator of myelopoiesis and atherogenesis. Arterioscler Thromb Vasc Biol. 2012;32(11):2547–52.

42. TeSlaa T, Ralser M, Fan J, Rabinowitz JD. The pentose phosphate pathway in health and disease. Nat Metab. 2023;5(8):1275–89.

43. Artyomov MN, Van den Bossche J. Immunometabolism in the Single-Cell Era. Cell Metabolism. 2020;32(5):710–25.

44. Gupta R, Gupta N. Pentose Phosphate Pathway. In: Gupta R, Gupta N, editors. Fundamentals of Bacterial Physiology and Metabolism. Singapore: Springer Singapore; 2021. p. 289–305.

45. Seo S-K, Kwon B. Immune regulation through tryptophan metabolism. Experimental & Molecular Medicine. 2023;55(7):1371–9.

46. Sorgdrager FJH, Naudé PJW, Kema IP, Nollen EA, Deyn PPD. Tryptophan Metabolism in Inflammaging: From Biomarker to Therapeutic Target. Frontiers in Immunology. 2019;10.

47. Chase TN, Murphy DL. Serotonin and central nervous system function. Annu Rev Pharmacol. 1973;13:181–97.

48. Herr N, Bode C, Duerschmied D. The Effects of Serotonin in Immune Cells. Frontiers in Cardiovascular Medicine. 2017;4.

49. Wu G, Bazer FW, Davis TA, Kim SW, Li P, Marc Rhoads J, et al. Arginine metabolism and nutrition in growth, health and disease. Amino Acids. 2009;37(1):153–68.

50. Lian J, Liang Y, Zhang H, Lan M, Ye Z, Lin B, et al. The role of polyamine metabolism in remodeling immune responses and blocking therapy within the tumor immune microenvironment. Frontiers in Immunology. 2022;13.

51. Ganeshan K, Chawla A. Metabolic regulation of immune responses. Annu Rev Immunol. 2014;32:609–34.

52. Codoñer-Franch P, Tavárez-Alonso S, Murria-Estal R, Herrera-Martín G, Alonso-Iglesias E. Polyamines Are Increased in Obese Children and Are Related to Markers of Oxidative/Nitrosative Stress and Angiogenesis. The Journal of Clinical Endocrinology & Metabolism. 2011;96(9):2821–5.

53. Ward ZJ, Bleich SN, Cradock AL, Barrett JL, Giles CM, Flax C, et al. Projected U.S. State-Level Prevalence of Adult Obesity and Severe Obesity. N Engl J Med. 2019;381(25):2440–50.

54. Lister NB, Baur LA, Felix JF, Hill AJ, Marcus C, Reinehr T, et al. Child and adolescent obesity. Nature Reviews Disease Primers. 2023;9(1):24.

55. Alfadhli EM. Maternal obesity influences birth weight more than gestational diabetes. BMC Pregnancy and Childbirth. 2021;21(1):111.

56. Sebire NJ, Jolly M, Harris JP, Wadsworth J, Joffe M, Beard RW, et al. Maternal obesity and pregnancy outcome: a study of 287,213 pregnancies in London. Int J Obes Relat Metab Disord. 2001;25(8):1175–82.

57. Ramsay JE, Ferrell WR, Crawford L, Wallace AM, Greer IA, Sattar N. Maternal obesity is associated with dysregulation of metabolic, vascular, and inflammatory pathways. J Clin Endocrinol Metab. 2002;87(9):4231–7.

58. Vrieling F, Stienstra R. Obesity and dysregulated innate immune responses: impact of micronutrient deficiencies. Trends in Immunology. 2023.

59. Zhou H-Y, Feng X, Wang L-W, Zhou R, Sun H, Chen X, et al. Bone marrow immune cells respond to fluctuating nutritional stress to constrain weight regain. Cell Metabolism.

60. Connor KM, Hsu Y, Aggarwal PK, Capone S, Colombo AR, Ramsingh G. Understanding metabolic changes in aging bone marrow. Experimental Hematology & Oncology. 2018;7(1):13.

61. Murphy MP. How mitochondria produce reactive oxygen species. Biochem J. 2009;417(1):1–13.

62. Giorgi C, Marchi S, Simoes ICM, Ren Z, Morciano G, Perrone M, et al. Mitochondria and Reactive Oxygen Species in Aging and Age-Related Diseases. Int Rev Cell Mol Biol. 2018;340:209–344.

63. Rubinsztein DC, Mariño G, Kroemer G. Autophagy and aging. Cell. 2011;146(5):682–95.

64. Houtkooper RH, Argmann C, Houten SM, Cantó C, Jeninga EH, Andreux PA, et al. The metabolic footprint of aging in mice. Sci Rep. 2011;1:134.

65. Mizushima N, Klionsky DJ. Protein turnover via autophagy: implications for metabolism. Annu Rev Nutr. 2007;27:19–40.

66. Muralimanoharan S, Gao X, Weintraub S, Myatt L, Maloyan A. Sexual dimorphism in activation of placental autophagy in obese women with evidence for fetal programming from a placenta-specific mouse model. Autophagy. 2016;12(5):752–69.

67. Liu X, Jiang X, Liu R, Wang L, Qian T, Zheng Y, et al. B cells expressing CD11b effectively inhibit CD4+ T-cell responses and ameliorate experimental autoimmune hepatitis in mice. Hepatology. 2015;62(5):1563–75.

68. Griffin DO, Rothstein TL. A small CD11b(+) human B1 cell subpopulation stimulates T cells and is expanded in lupus. J Exp Med. 2011;208(13):2591–8.

69. Chen EP, Markosyan N, Connolly E, Lawson JA, Li X, Grant GR, et al. Myeloid Cell COX-2 deletion reduces mammary tumor growth through enhanced cytotoxic T-lymphocyte function. Carcinogenesis. 2014;35(8):1788–97.

70. Eruslanov E, Daurkin I, Ortiz J, Vieweg J, Kusmartsev S. Pivotal Advance: Tumor-mediated induction of myeloid-derived suppressor cells and M2-polarized macrophages by altering intracellular PGE₂ catabolism in myeloid cells. J Leukoc Biol. 2010;88(5):839–48.

71. Na YR, Yoon YN, Son DI, Seok SH. Cyclooxygenase-2 inhibition blocks M2 macrophage differentiation and suppresses metastasis in murine breast cancer model. PLoS One. 2013;8(5):e63451.

72. Cuccurullo C, Fazia ML, Mezzetti A, Cipollone F. COX-2 expression in atherosclerosis: the good, the bad or the ugly? Curr Med Chem. 2007;14(15):1595–605.

73. Markosyan N, Chen EP, Evans RA, Ndong V, Vonderheide RH, Smyth EM. Mammary carcinoma cell derived cyclooxygenase 2 suppresses tumor immune surveillance by enhancing intratumoral immune checkpoint activity. Breast Cancer Res. 2013;15(5):R75.

74. Markosyan N, Chen EP, Ndong VN, Yao Y, Sterner CJ, Chodosh LA, et al. Deletion of cyclooxygenase 2 in mouse mammary epithelial cells delays breast cancer onset through augmentation of type 1 immune responses in tumors. Carcinogenesis. 2011;32(10):1441–9.

