## Supplemental Table 1 for "Metabolic abnormalities in the bone marrow cells of young offspring born to obese mothers"

| Target | Fluorochrome | Clone | Company | Cat # |
| --- | --- | --- | --- | --- |
| CD45 | BUV395 Rat Anti-Mouse | 30-F11 | BD Biosciences | 564279 |
| CD4+ | BUV615 Rat Anti-Mouse | RM4-5 | BD Biosciences | 751486 |
| CD19+ | BUV661 Rat Anti-Mouse | 1D3 | BD Biosciences | 612971 |
| CD3 Molecular Complex | BUV737 Rat Anti-Mouse CD3 | 17A2 | BD Biosciences | 612803 |
| CD8+ | BUV805 Rat Anti-Mouse CD8a | 53-6.7 | BD Biosciences | 612898 |
| CX3CR1 | Brilliant Violet 421™ | CXCR3-173 | BioLegend | 126521 |
| MHCII | eFluor™ 450 | M5/114.15.2 | Invitrogen | 48-5321-82 |
| TNFα | Brilliant Violet 510™ anti-mouse | MP6-XT22 | BioLegend | 506339 |
| CD11c | Brilliant Violet 605™ anti-mouse | N418 | BioLegend | 117334 |
| CD86 | Brilliant Violet 650™ anti-mouse | GL-1 | BioLegend | 105035 |
| PD-L1 | Brilliant Violet 711™ anti-mouse | 10F.9G2 | BioLegend | 124319 |
| CD103 | BD OptiBuild™ BV750 Rat Anti-Mouse | Ber-ACT8 | BD Biosciences | 747478 |
| F4/80 | Brilliant Violet 785™ anti-mouse F4/80 Antibody | BM8 | BioLegend | 123141 |
| COX-2 | Alexa Fluor™ 488,  anti-mouse/human | COX 229 | Invitrogen (TF) | 35-8200 |
| Ly-6C | PerCP/Cyanine5.5 anti-mouse | HK1.4 | BioLegend | 128012 |
| IFNγ | PE, rat anti mouse/human | XMG1.2 | eBioscience (TF) | 12-7311-82 |
| CD206 | PE/Cyanine7 anti-mouse | C068C2 | BioLegend | 141720 |
| ApoE | Alexa Fluor® 647 anti-mouse | 1G5G9 | Novus Biologicals | NB110-55454AF647 |
| CD11b | Alexa Fluor® 700 anti-mouse/human | M1/70 | BioLegend | 101222 |
| CD38 | APC/Fire™ 810 anti-mouse | 90 | BioLegend | 102746 |
| Ly-6G | APC/Fire™ 750 anti-mouse | 1A8 | BioLegend | 127651 |

**Supplemental Table 1**. Flow cytometry antibody panel.
