## Supplemental Table 2 for "Metabolic abnormalities in the bone marrow cells of young offspring born to obese mothers"

| **Gene** | **Forward (5’-3’)** | **Reverse (5’-3’)** |
| --- | --- | --- |
| ***Il6*** | GTT CTC TGG GAA ATC GTG GA | TCC AGT TTG GTA GCA TCC ATC |
| ***Tnf*** | TGC TCT GTG AAG GGA ATG GG | ACC CTG AGC CAT AAT CCC CT |
| ***Ifng*** | CAG CAA CAG CAA GGC GAA AAA GG | TTT CCG CTT CCT GAG GCT GGA T |
| ***ODC*** | CCTTGTGAGGAGCTGGTGATA | GGTCCAGAATGTCCTTAGCAGT |
| ***ACTB*** | AAG GCC AAC CGT GAA AAG AT | GTG GTA CGA CCA GAG GCA TAC |

**Supplemental Table 2**. Nucleotide sequences of RT-PCR primers.
