## Supplementary figures and images for "Metabolic abnormalities in the bone marrow cells of young offspring born to obese mothers"

### Supplemental Figure 1

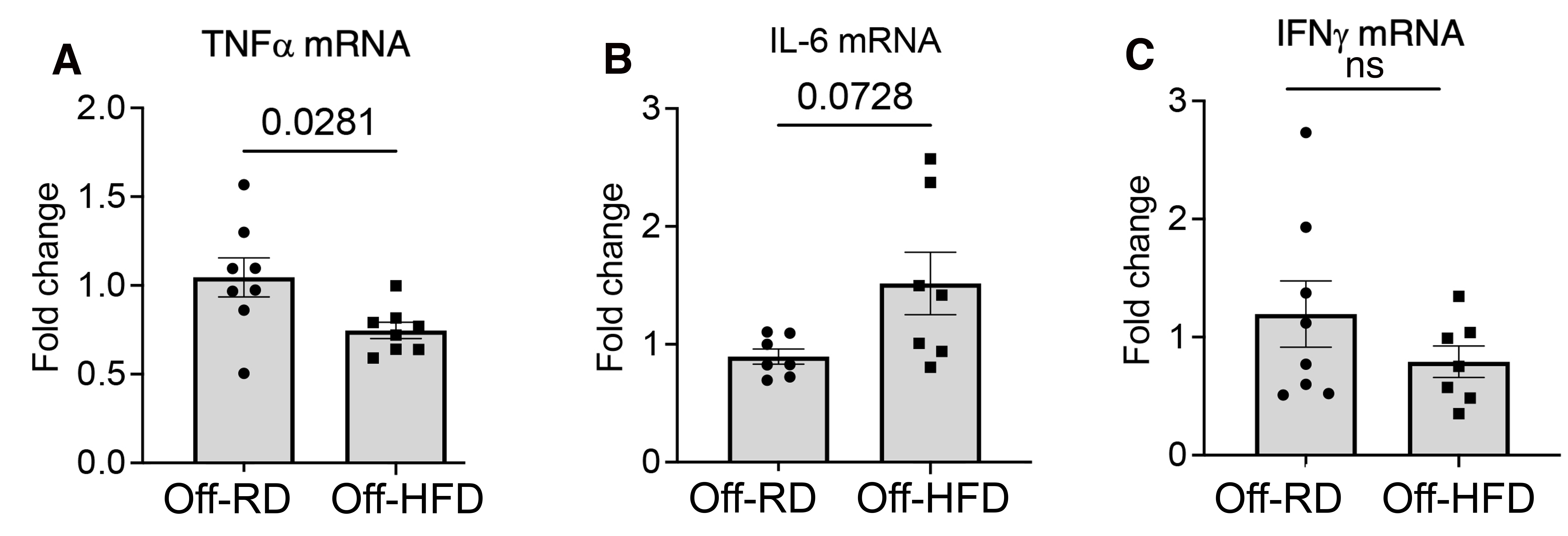
